# Mammary gland metabolism and its relevance to the fetoplacental expression of cytokine signaling in Caveolin-1 null mice

**DOI:** 10.1101/2025.04.13.648647

**Authors:** Shankar P. Poudel, Maliha Islam, Thomas B. McFadden, Susanta K. Behura

## Abstract

Mice lacking Caveolin-1 (*Cav1*), a major protein of the lipid raft of plasma membrane, show dysregulated cellular proliferation of mammary gland and an abnormal fetoplacental communication during pregnancy. The aim of this study is to better understand the functional links of mammary gland metabolism with gene expression of the placenta and fetus. Untargeted metabolomics analysis was performed to examine changes in mammary gland metabolism due to the absence of *Cav1*. Integrative metabolomics and transcriptomics analyses were applied to untangle functional links of metabolic pathways of the mammary gland with the gene expression changes of the placenta and fetus. The findings of this study show that metabolism and gene expression of the mammary gland are significantly impacted due to the loss of *Cav1*. Genes associated with specific metabolic and signaling pathways show coordinated expression changed in the placenta, mammary gland and fetal brain in *Cav1*-null mice. The cytokine signaling pathway emerges as a key player of the molecular crosstalk among the mammary gland, placenta and fetal brain. By interrogating the single-nuclei gene expression data of placenta and fetal brain previously generated from *Cav1*-null mice, the study further reveals that these metabolic and signaling genes are differentially regulated in specific cell types of the placenta and fetal brain. The findings of this study expand our understanding about the role of mammary gland metabolism in the regulation of fetoplacental communication in mammalian pregnancy.

## Introduction

The growth of mammary gland and placenta occur in a correlated manner during mammalian pregnancy (Nagasawa 1982). The placenta-derived lactogen promotes the growth of mammary gland to prepare for lactation (Wrenn et al. 1966). In mice, two types of placental lactogen (PL-I and PL-II) are produced by the placenta, mostly by the trophoblast giant cells. The PL-I is synthesized immediately after the embryo is implanted whereas the PL-II is synthesized during the mid-gestation period, but both are required to maintain the pregnancy and promote mammary gland growth (Soslow et al. 2006). On gestation day 15 (d15), the mouse placenta becomes fully developed and functional (Panja and Paria 2021) after which the mammary gland grows rapidly (Desjardinus et al. 1968) along with the rapid growth of the fetus (Woods et al. 2018).

The mammary gland development is mediated by binding of the pituitary hormone prolactin (Prl) (Napso et al. 2018) to its receptor which then activates the *Jak-2*/*Stat5a* signaling pathway. The Caveolin-1 (*Cav1*), a plasma member protein, functions as negative regulator of Jak-2 tyrosine kinase through the inhibition of prolactin induced *Stat5a* tyrosine phosphorylation and DNA binding activity. The *Cav1*-null mice show premature growth of the lobuloalveolar compartment of the mammary gland (Park et al. 2002). These mice also exhibit premature aging and neurodegeneration at an early adult life (Head et al. 2010). Moreover, *Cav1* also plays other key roles in cellular metabolism, cell proliferation and apoptosis, trophoblast invasion, angiogenesis, intracellular and extracellular transport and signaling (Feng et al. 2012; Frank et al. 2006; J. Li et al. 2005; Volonte and Galbiati 2020). *Cav1* down-regulation promotes mitogenic activity in mammary cell division (Zhang et al. 2005). Also, *Cav1* functions as a tumor suppressor gene and its knockout leads to hyperproliferative phenotype (Razani et al. 2001). In human, overexpression of *Cav1* is associated with the metaplastic breast carcinomas (Savage et al. 2007). In addition, *Cav1* plays an important role in the regulation of immune cells (Wang et al. 2023). The *Cav1*-null mice show an increased activity of NF-κB signaling and inflammatory fibrosis in heart tissues (Gong et al. 2023). *Cav1* influences regulation of inflammatory response through *Stat3*/ *NF-*κ*B* pathway, and its ablation leads to compromised immune response to inflammation (Yuan et al. 2011). Also, *Cav1*-null mice show a higher level of cytokine production in the lymphocytes than control mice suggesting importance of *Cav1* in the regulation of immune response to inflammation (Codrici et al. 2018). While cytokine signaling plays important roles in the fetoplacental communication in mammalian pregnancy (Hauguel-de Mouzon and Guerre-Millo 2006; Yockey and Iwasaki 2018), it is unknown if loss of *Cav1* alters cytokine signaling to influence the mammary gland and fetoplacental communication.

Earlier we showed that lack of *Cav1* deregulates fetal brain metabolism (Islam and Behura 2023), and leads to an abnormal fetoplacental crosstalk during pregnancy (Islam and Behura 2024). In addition, by performing single-nuclei RNA sequencing (snRNA-seq), we have further shown that specific cell types of mouse placenta and fetal brain express genes in a coordinated manner due to the lack of *Cav1* (Islam and Behura 2024). In the present study, we hypothesize that deregulation of mammary gland growth influences the fetoplacental regulation in the *Cav1*-null mice. To test this hypothesis, the objectives of the present study are to perform integrative analysis of metabolomics and gene expression data to untangle the functional links of mammary gland with the regulation of placenta and fetal brain in *Cav1*-null mice.

## Experimental

### Mice breeding and sample collection

The B6.Cg-*Cav1^tm1Mls^*/J (*Cav1*-null) and control (C57BL/6J) mice were purchased from the Jackson Laboratory (stock numbers: 000664 and 007083 respectively). Adult male and female mice (7 weeks old) were paired in cages to induce pregnancy. The females were checked for vaginal plug. The start of pregnancy (day 1) was considered when a vaginal plug was observed. The control and *Cav1*-null pregnant mice were euthanized in triplicates on gestation day 15, and the mammary gland samples were dissected from mice of both the groups (n=3 each). Along with the mammary gland samples, the placentae were also collected from the *Cav1*-null mice. The samples were washed in sterile phosphate buffer saline, snap frozen in liquid nitrogen and stored in -80°C for downstream analyses. The animal procedures were approved by the Institutional Animal Care and Use committee of the University of Missouri, and were conducted according to the guide for Care and Use of Laboratory Animals (National Institute of Health, USA).

### Metabolomics analysis

The mammary gland samples, in triplicates from the control and *Cav1*-null mice, were subjected to an untargeted metabolomics analysis by gas chromatography mass spectrometry (GC-MS). The samples were processed at the University of Missouri Metabolomics Center as described in our earlier studies (Islam and Behura 2023; Poudel and Behura 2024). An Agilent 6890 GC coupled to a 5973N MSD mass spectrometer was used for GC-MS analysis. The spectral analysis was performed by the AMDIS (Automate Mass-spectral Deconvolution and Identification System) (Meyer et al. 2010), and metabolites were identified using a commercial NIST17 mass spectral library. The abundance of the metabolites was determined by the Metabolomics Ion-Based Data Extraction Algorithm (MET-IDEA) (Broeckling et al. 2006). The metabolomics data was processed using the *MetaboAnalyst* tool (Xia et al. 2009). Differential metabolite analysis was performed using the R package *MetaboDiff* (Mock et al. 2018). The raw and processed data of the metabolomics assay are available at Metabolomics Workbench (Sud et al. 2016) under study ID ST003855 and accession ID SbsM9645.

### RNA-seq analysis

Total RNA was isolated from the mammary gland (in triplicates) of control and *Cav1*-null mice using AllPrep DNA/RNA Mini Kit (Qiagen, Cat No./ID:80204) following the manufacturers’ instructions. RNA was also isolated from placenta (also in triplicates) of three *Cav1*-null mice. RNA integrity was determined using an Agilent 2100 bioanalyzer. Total RNA was used for preparation of libraries followed by library sequencing (RNA-seq) by the University of Missouri Genomics Technology facility. Each library was sequenced to 20 million paired end reads of 150 bases using an Illumina NovaSeq 6000 sequencer. The quality of raw sequences were checked with *FastQC* followed by trimming adaptors and base quality trimming using *fastp* (Chen et al. 2018). The reads were than mapped to the mouse reference genome *GRCm39* using the *subread* aligner (Liao et al. 2013). The number of reads mapped to the individual genes was determined using the *featureCounts* tool (Liao et al. 2014). The raw and processed data have been submitted to the Gene Expression Omnibus (GEO) database under the accession number GSE268570. The gene expression data of mammary gland from *Cav1*-null and control mice was compared with the gene expression data of placenta and fetal brain. The gene expression data of the fetal brain (GSE215139) of *Cav1*-null and control mice were generated from our earlier work (Islam and Behura 2023). The gene expression data of the day-15 placenta of C57BL/6J mice (accession GSE215139 was obtained from another study by our lab (Dhakal et al. 2021). Differential gene expression analysis was performed between the mammary gland and placenta samples using the R package *edgeR* (Robinson et al. 2010). The Bioconductor mouse genome annotation database (*org.Mm.eg.db*) was used to identify the Kyoto Encyclopedia of Genes and Genomes (KEGG) metabolic genes associated with the differentially expressed genes.

### Pathway enrichment analysis

The pathway enrichment analysis was performed using the PANTHER tool (Mi et al. 2013). In this analysis, the significance of enrichment was assessed by the Fisher exact test followed by multiple corrections of raw *p*-values to False Discovery Rate (FDR).

### Hierarchical clustering

The genes associated with the enriched pathways were analyzed by hierarchical clustering of gene expression variation among the mammary gland, placenta and fetal brain of the control and *Cav1*-null mice using the *R* package *dendextend* (Galili 2015). The Euclidean distances of gene expression variation were clustered by ward.D2 as the method of agglomeration. Comparison between cluster dendrograms between mammary gland, placenta and fetal brain were performed using *dendextend*.

### Network and key player analysis

The gene regulatory network analysis was performed using *minet* (Meyer et al. 2008). The mutual information (MI) measures (Steuer et al. 2002) of gene expression changes were used for inferring networks. Gene sets representing the different functions (metabolic and signaling pathways) were used for MI calculation in a pair-wise manner between genes followed by developing the weighted adjacency matrix by the maximum relevance minimum redundancy (MRMR) method (Radovic et al. 2017). Then, mutual information network analysis was performed using *minet* (Meyer et al. 2008). The key players of the inferred networks were predicted by centrality test (Koschützki and Schreiber 2008) using the R package ‘*key player*’.

## Results

### Mammary gland metabolism in Cav1-null mice

We performed an untargeted metabolomics analysis of day-15 mammary gland from the control and *Cav1*-null mice to investigate if loss of *Cav1* impacted mammary gland metabolism during pregnancy. The gestation day 15 was chosen because placenta becomes fully formed at this time point (Panja and Paria 2021). The metabolomics analysis detected a total of 141 metabolites which are provided in **Supplementary Table 1**. The heatmap in **Figure 1A** shows the pattern of variation of the metabolites. The mutual information analysis, which is a method to quantify the non-linear relationships between variables (Steuer et al. 2002), revealed the relationships of metabolic changes among the mammary glands (**Figure 1B**). We identified metabolites (n=81) that showed differential rank order in the level of expression in the mammary glands of *Cav1*-null compared to control mice (**Supplementary Table 2**). Differential metabolomics analysis (Mock et al. 2018) identified specific metabolites that were significantly changed in the mammary gland due to the absence of *Cav1* (**Table 1**).

**Figure 1.**
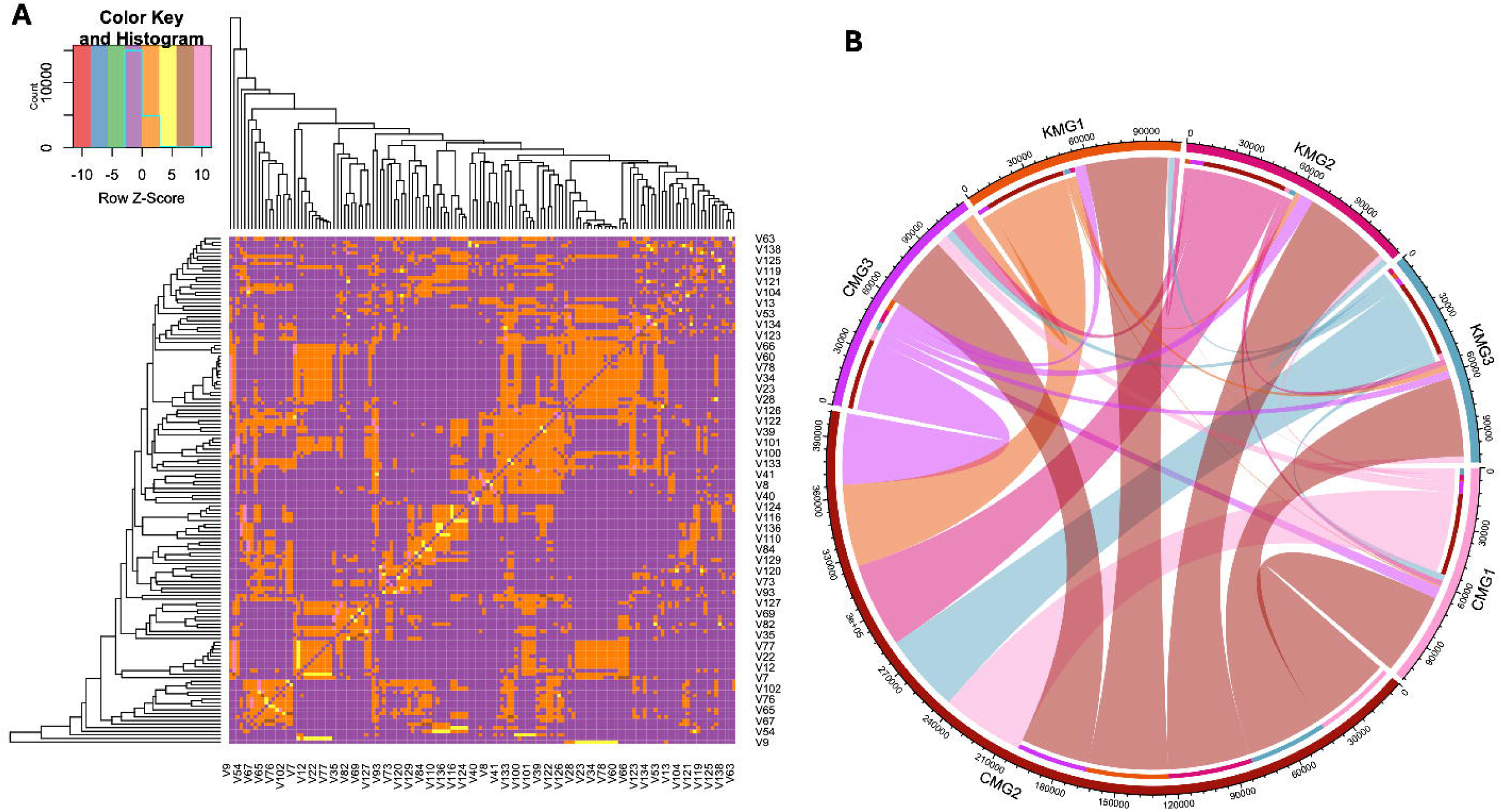
Metabolic variation in mammary gland of *Cav1*-null relative to control mice. **A**. Heat map showing relationship of metabolites, in a pairwise manner, based on their variation in mammary gland of *Cav1*-null and control mice. The rows and columns show the individual metabolites. They are labeled as v1, v2, etc. instead of the metabolite names. The color scale shows the z-value of metabolic variation calculated from pairwise mutation information (MI) distance matrix. **B**. Circos plot showing the relationship among the individual mammary gland (MG) samples of control (C) and *Cav1* knockout (K) mice (n=3). The samples are shown on the circumference. The arcs (color coded) connecting the samples represent the magnitude of MI between the two samples.

**Table 1.** Specific metabolites significantly changed in the mammary gland of *Cav1*-null relative.

| Metabolite | Change in level<br>(relative to control) | Fold change | Adjusted P-value (FDR) |
| --- | --- | --- | --- |
| L-Threonine | Increase | 2.458 | 0.012 |
| L-Glutamic Acid | Increase | 9.165 | 0.012 |
| L-Lysine | Increase | 14.15 | 0.015 |
| L-Proline | Increase | 5.068 | 0.019 |
| L-Phenylalanine | Increase | 7.982 | 0.021 |
| Succinic Acid | Increase | 2.372 | 0.038 |
| Glyceric acid-3-phosphate | Increase | 3.236 | 0.038 |
| L-Tyrosine, O-trimethylsilyl-,<br>trimethylsilyl ester | Increase | 7.012 | 0.039 |
| Urea | Increase | 2.046 | 0.039 |
| L-Aspartic acid | Increase | 2.229 | 0.039 |
| Pentasiloxane Dodecamethyl | Decrease | 0.324 | 0.005 |

### Expression changes of metabolic genes in mammary gland of Cav1-null mice

RNA sequencing (RNA-seq) was performed to analyze gene expression of mammary gland of *Cav1*-null and control day-15 pregnant mice. The gene expression data was analyzed in an integrative manner with the metabolomics data by applying the approach described earlier (Islam and Behura 2023). In this approach, the genes identified from the RNA-seq data were mapped to the KEGG (Kyoto Encyclopedia of Genes and Genomes) pathways (Kanehisa et al. 2017). The mapping results identified a set of 429 genes related to different metabolic pathways of mouse (**Supplementary Table 3**). Next, the expression of these genes in the mammary gland was compared with the expression of those genes in the placenta and fetal brain of *Cav1*-null and control mice (**Figure 2**). The comparative analysis identified a set of 76 metabolic genes (**Supplementary Table 4**) that were upregulated in the mammary gland but downregulated in the placenta and fetal brain due to the absence of *Cav1*.

**Figure 2.**
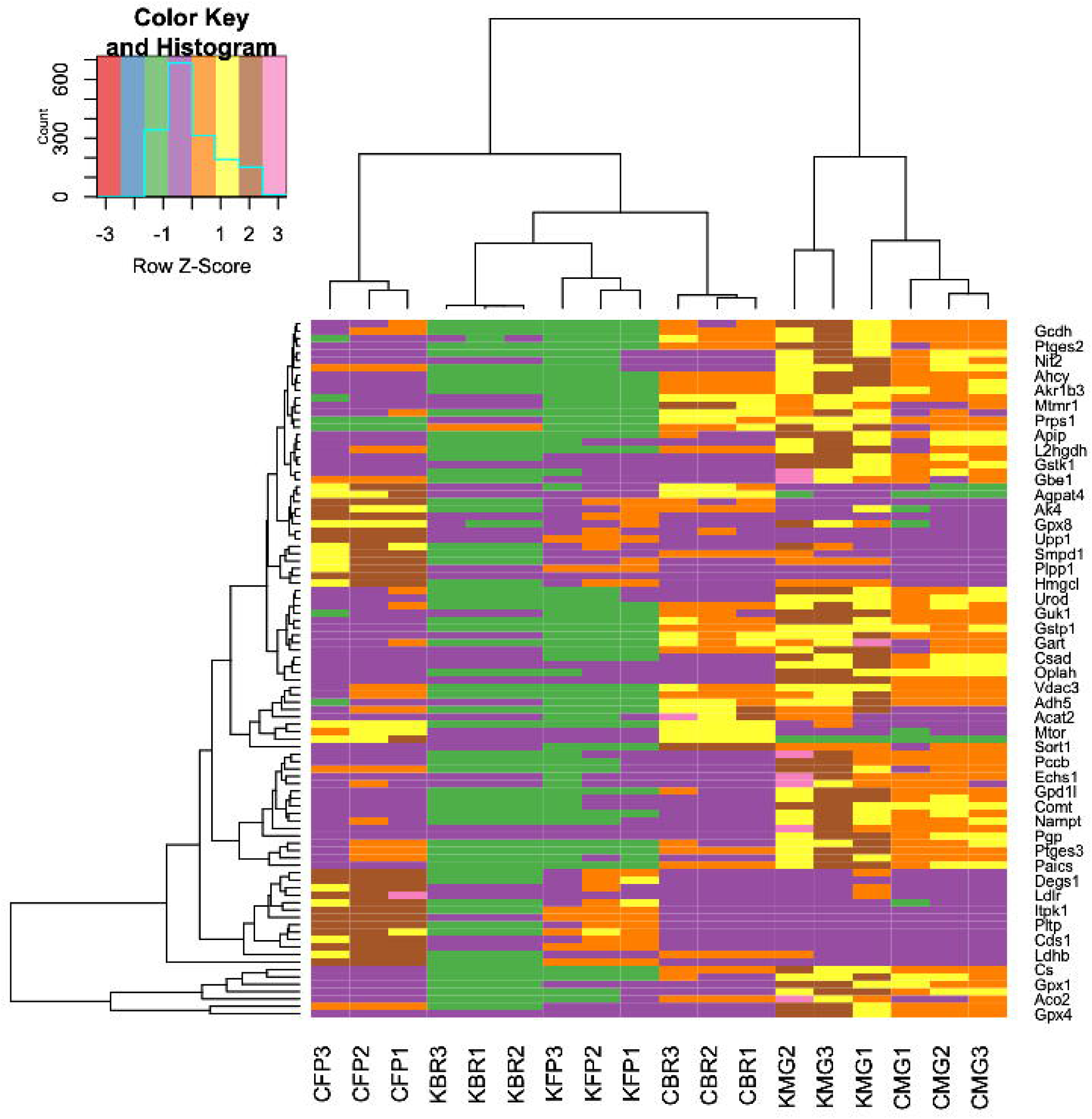
Expression changes of metabolic genes in the mammary gland, placenta and fetal brain. The heatmap shows expression variation of metabolic genes among the mammary gland (MG), placenta (FP) and fetal brain (BR) of control (C) and *Cav1*-null (K) mice (n=3). The genes are shown in rows and samples are shown in columns. The color code of gene expression variation is according to the z-score scale shown in the top left.

### Expression clusters of metabolic and signaling genes in mammary gland

Hierarchical clustering (Ward Jr. 1963) of gene expression clusters among the mammary gland, placenta and fetal brain samples identified 9 clusters. The genes associated with the individual clusters are listed in **Supplementary Table 5**. Different number of metabolic genes were associated with these clusters. The odds ratio and significance of association of metabolic genes with the individual clusters are shown in **Table 2**. Clusters 4 and 7 showed a significant enrichment with the metabolic genes. Moreover, these clusters also showed significant enrichment for specific pathways based on PANTHER pathway enrichment analysis (Mi et al. 2013). They included apoptosis signaling pathway, EGF receptor signaling pathway, toll receptor signaling pathway, integrin signaling pathway, and PDGF signaling pathway among others (**Supplementary Table 6)**. Being associated with the same clusters, the findings suggest that the metabolic and signaling genes are coordinately regulated among the mammary gland, placenta and fetal brain in response to the absence of *Cav1*. We referred this to as the mammary gland-placenta-fetal brain (MBP) axis.

**Table 2.** Odds ratio and significance of association of metabolic genes with different gene expression clusters. CI: confidence interval (upper and lower values). The significant clusters and associated P-values are shown in bold. n.a: not applicable.

| Cluster # | Metabolic genes | Non-metabolic genes | Odds ratio | CI (95%) | P-value |
| --- | --- | --- | --- | --- | --- |
| 1 | 0 | 394 | n.a | n.a | n.a |
| 2 | 0 | 79 | n.a | n.a | n.a |
| 3 | 4 | 824 | 0.508 | 0.135, 1.35 | 0.249 |
| **4** | 26 | 1392 | 1.929 | 1.194, 3.018 | **0.005** |
| 5 | 13 | 1080 | 1.251 | 0.64, 2.255 | 0.418 |
| 6 | 40 | 5515 | 0.757 | 0.509, 1.11 | 0.148 |
| **7** | 9 | 339 | 2.721 | 1.196, 5.45 | **0.009** |
| 8 | 2 | 158 | 1.315 | 0.155, 4.964 | 0.667 |
| 9 | 0 | 18 | n.a | n.a | n.a |

### Cytokine signaling as a key player of the regulation of the MPB axis

To further understand the regulation of MPB axis due to the loss of *Cav1*, we applied an integrative method that measured the distance between gene expression cluster of the metabolic and signaling genes (Galili 2015), and then inferred their crosstalk pattern by neural network analysis (Meyer et al. 2008). In this analysis, we used the R package *dendextend* (Galili 2015) to generate dendrograms of the metabolic and signaling genes (**Figure 3)**. The distance between these clusters was measured in a pair-wise manner to predict their crosstalk pattern by mutual informatic network analysis (Meyer et al. 2008) (**Figure 4)**. Centrality test (Koschützki and Schreiber 2008) of the inferred network further showed that the cytokine signaling was a key player of the observed network suggesting that cytokine signaling played a central role in the regulation of the MPB axis due to the absence of *Cav1*.

**Figure 3.**
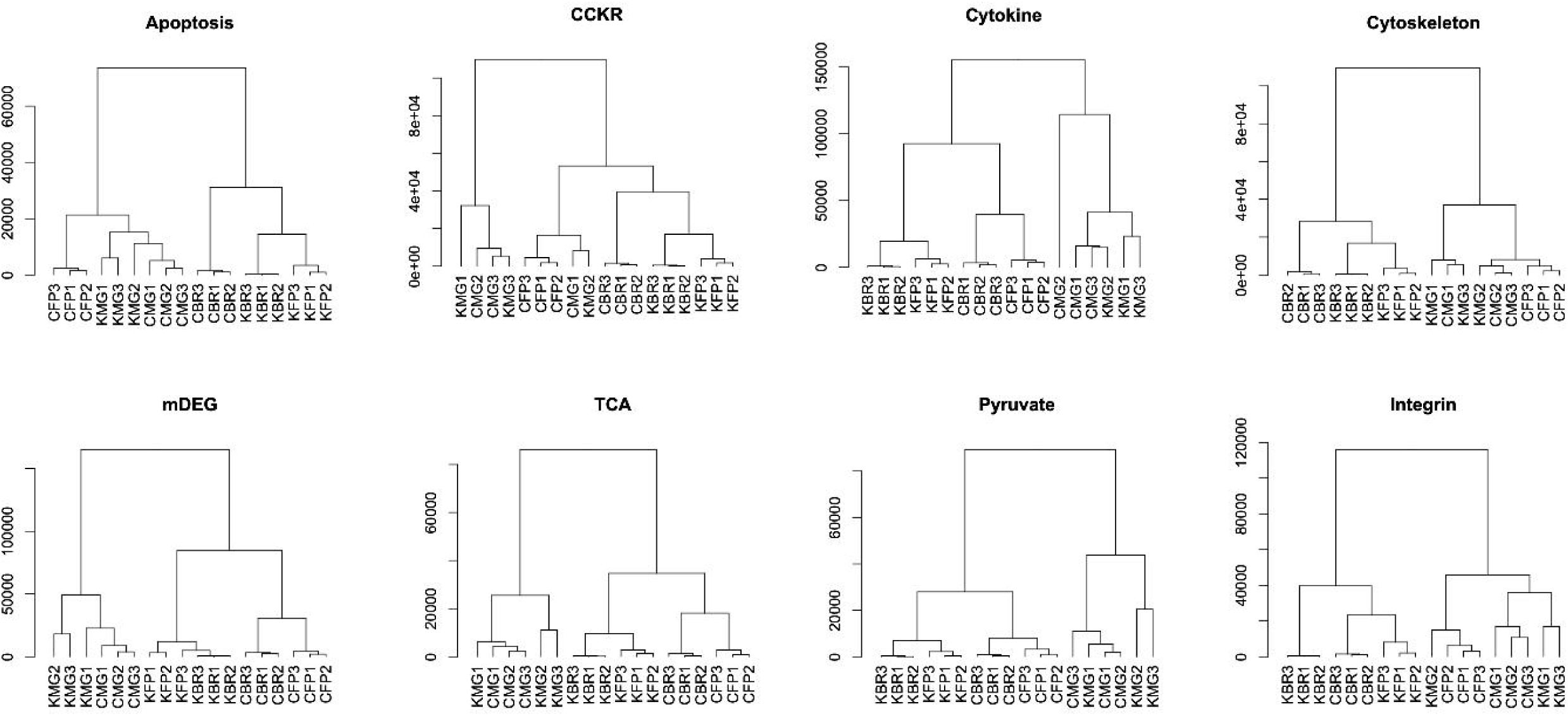
Hierarchical clusters of metabolic and signaling genes. The samples are shown as mammary gland (MG), placenta (FP) and fetal brain (BR) of control (C) and *Cav1*-null (K) mice (n=3). The cluster relationships among the samples are shown for each gene type which is indicated on the top of the dendrogram plot. The scale shown in each plot represents the branch length of the dendrogram.

**Figure 4.**
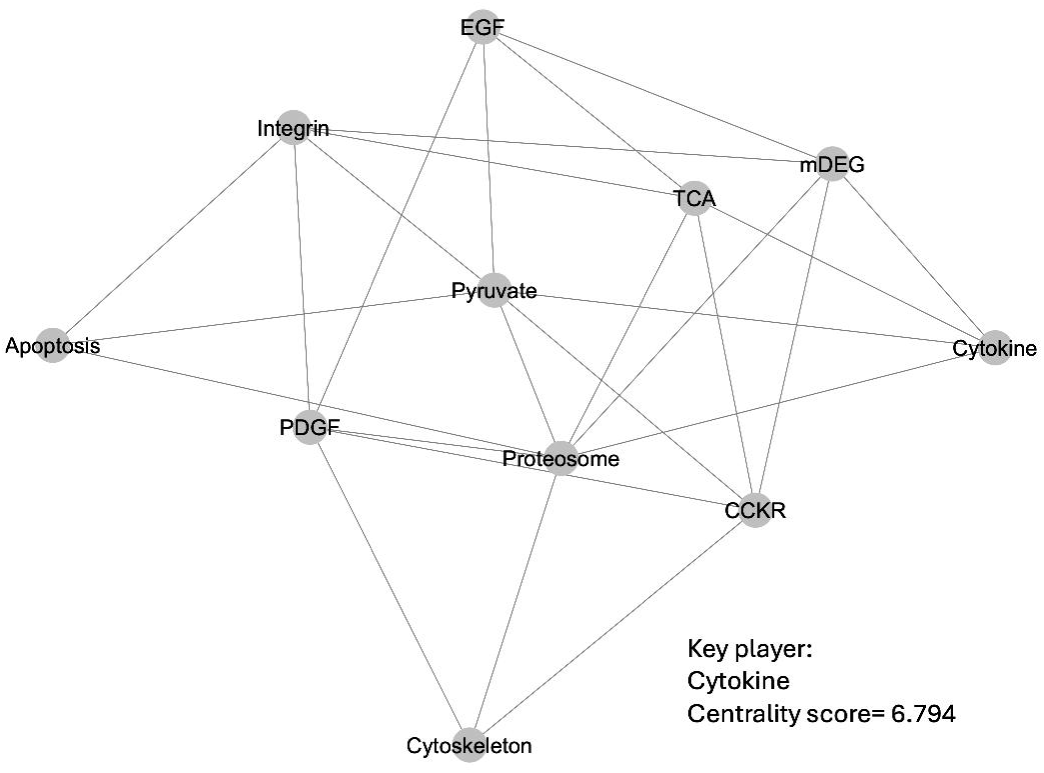
Crosstalk of metabolic and signaling pathways. The plot shows the mutual information network pattern of different metabolic and signaling pathways based on expression variation of the pathway genes. The nodes represent the pathways, and the edges represent the crosstalk pattern among them. The mDEGs, shown as one of the nodes, represent the metabolic genes that were upregulated in the mammary gland but downregulated in the placenta and fetal brain in *Cav1*-null mice (listed in Supplementary Table 4). The centrality score of the node representing cytokine signaling is shown.

### Expression of the metabolic and signaling genes in placenta and fetal brain cells

We further wanted to know how the metabolic and signaling genes were regulated in the placenta and fetal brain cells. Towards this objective, we extracted the single-nuclei RNA-seq data of placenta and fetal brain cells generated from *Cav1*-null and control mice from our earlier studies (Islam and Behura 2023, 2024). These data are publicly available in Gene Expression Omnibus database under the accession numbers GSE214759 and GSE248126. The heatmaps in **Figure 5** show the expression patterns of the metabolic and the signaling genes (see Supplementary Table 4) in the individual cell types of the fetal brain and placenta of the control and *Cav1*-null mice. The heatmaps show that these genes are expressed at a higher level in the cytotrophoblast cells (CTB), which are stem cells from which the synsytiotrophoblast (STB) cells are differentiated, than any other cell types of the placenta. While the extravillious trophoblast (EVT) and STB cells were closely related in expression of these genes in the control mice, they were distantly related in the *Cav1*-null mice. We calculated the aggregated expression of the individual genes in the placenta and brain cells (**Supplementary Table 7)**, and used the data to performed Principal Component Analysis (PCA). The results of this analysis are shown in **Figure 6**. The percentage of gene expression variation explained by the two principal axes (PC1 and PC2) are shown in this figure. The percentage of data variation decreased in PC1 but increased in PC2 for the fetal brain due to the absence of *Cav1*. In the placenta, the data variation increased in PC1 but decreased in PC2 in *Cav1*-null compared to control mice. The signal transducer and activator of transcription 5A (Stat5a), which is activated by cytokines to influence the growth of mammary gland of *Cav1*-null mice (Park et al. 2002), showed a suppressed expression pattern in the EVT cells of placenta as well the radial glial cells of fetal brain due to the absence of *Cav1* (**Supplementary Figure 1**) further supporting an impact of *Cav1* ablation in the regulation of MPB axis in mice.

**Figure 5.**
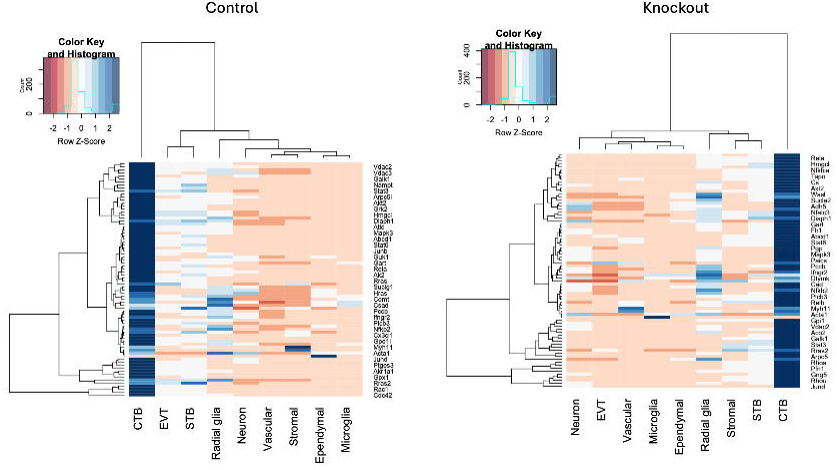
Expression variation of metabolic, and signaling genes in different cell types of the placenta and fetal brain. The heatmap on the left shows the expression patterns of these genes in the control mice. The heatmap on the right shows the expression patterns of these genes in the *Cav1*-null mice. The columns represent the cell types of the placenta and fetal brain. The rows represent the genes. The heatmaps are generated based on the aggregated expression of the gene for each cell type. The color code of gene expression variation is according to the z-score scale shown in the top left. CTB: cytotrophoblasts, EVT: extravillious trophoblasts, and STB: syncytiotrophoblasts.

**Figure 6.**
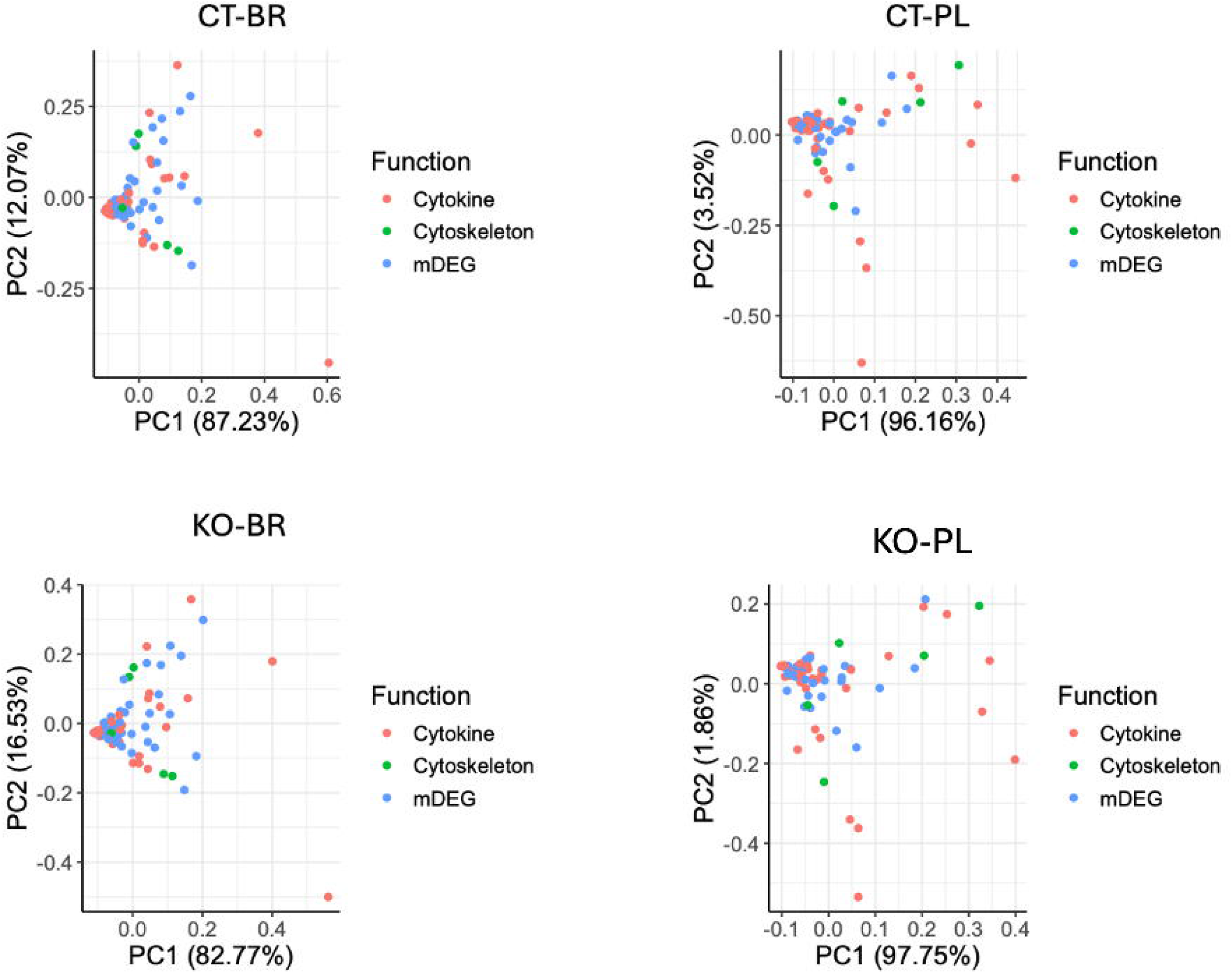
Expression variation of genes related to metabolic and signaling genes. The principal component analysis (PCA) plots show the percentage of gene expression variation by the major axes (PC1 and PC2). The color-coded dots represent the three categories of genes listed on the right of each plot. The mDEGs, shown as one of the gene categories, represent the metabolic genes that were upregulated in the mammary gland but downregulated in the placenta and fetal brain in *Cav1*-null mice (listed in Supplementary Table 4). The plots on the top represent gene expression variation in the fetal brain and placenta of control mice (CT-BR and CT-PL). The plots on the bottom represent gene expression variation in the fetal brain and placenta of *Cav1* knockout mice (KO-BR and KO-PL).

## Discussion

This study investigated metabolic changes of mammary gland and explored their relationship with the regulation of signaling genes in the placenta and fetal brain in mice. In this study, we used *Cav1*-null mice that show abnormal growth of mammary gland during pregnancy (Park et al. 2002). Untargeted metabolomics analysis identified metabolites that changed in the mammary gland due to the absence of *Cav1*. They included specific amino acids such as lysine that controls mammary gland development through PI3K/AKT/mTOR signaling pathway (W. Li et al. 2022), and glutamic acid that is required for synthesis of the milk protein casein (Barry 1956), The succinic acid, an intermediate in the pyruvate metabolism that plays key roles in energy production and the synthesis of milk components like lactose and fatty acids (Mazurek et al. 1999) was also increased in the mammary gland due to the absence of *Cav1*. By comparing the metabolites of mammary gland from the present study with the metabolites in the fetal brain of *Cav1*-null mice (Islam and Behura 2023), we observed that the proline level increased in the mammary gland but decreased in the fetal brain due to the absence of *Cav1*. The urea level increased in the mammary gland as well as fetal brain in *Cav1*-null mice. Increase in brain urea level is linked to Alzheimer’s disease in humans (Hansmannel et al. 2010; Ju et al. 2022), and *Cav1*-null mice show neurodegeneration patterns similar to that of human Alzheimer’s disease but at an early adult life (Head et al. 2010).

The integrated metabolomics and transcriptomics analysis showed that several metabolic genes changed in a coordinated manner in the mammary gland relative to the placenta and fetus. They were significantly (*p* < 0.05) upregulated in the mammary gland but downregulated in the placenta and fetal brain of the *Cav1*-null mice (**Supplementary Table 4**). Pathway enrichment analysis that specific metabolic and signaling pathways were significantly enriched with these genes (**Supplementary Table 5**). The pyruvate metabolic pathway was one of these. Expression of pyruvate metabolic genes increased in the mammary gland but decreased in the placenta and fetal brain of *Cav1*-null mice. Pyruvate metabolism plays important roles in the mammary gland growth (Raafat et al. 1963). The deregulation of pyruvate metabolism is also associated with neurodegeneration (Gray et al. 2013) which is observed in *Cav1*-null mice (Head et al. 2010). As *Cav1*-null pregnant mice show accelerated growth of the mammary gland (Park et al. 2002), this finding suggests that pyruvate metabolism of mammary gland may play a role in the regulation of MBP axis to influence the placenta and subsequently the fetal brain. This supports the idea that metabolism of mammary gland can affect pregnancy and fetal programming (Anhê and Bordin 2022; Bazer et al. 2004). We further observed genes related to specific metabolic and signaling pathways were expressed in a coordinated manner as evident from network analysis (**Figure 4**). It is known that functional links between metabolism with and signal transduction can influence growth and developmental processes (Chandel 2021). Moreover, metabolites act as sensor to modulate signal transduction networks that allows cells to coordinate the growth and development (Ward and Thompson 2012). By performing network analysis and centrality tests (Koschützki and Schreiber 2008; Meyer et al. 2008), we further observed that cytokine signaling acted as a key player of the metabolism-signaling network in the mammary gland, placenta and fetal brain (**Figure 4**). This suggested that metabolic changes in the mammary gland is functionally linked to the transcriptional crosstalk of signaling genes of the MPB axis.

Furthermore, genes associated with the MPB axis showed differential expression patterns in specific cell types of the placenta and fetal brain (**Figure 5**). The data in **Figure 5** shows that the cytotrophoblast cells of the placenta and the radial glia cells of the fetal brain expressed the metabolic and signaling genes of MPB axis at a higher level than all other cell types. The cytotrophoblasts are the stem cells of placenta from which STB and EVT cells are differentiated (Kolahi et al. 2017), and the radial glia cells are the neural stem cells of fetal brain (Malatesta et al. 2008). Our finding is consistent with the idea that stem cells are associated with a higher level of metabolism to support the signal transduction and differentiation processes of these cells (Jackson and Finley 2024; Shyh-Chang and Ng 2017). Our data further showed that the EVT cells of placenta were associated with a diminished expression of *Stat5a* in the absence of *Cav1* (**Supplementary Figure 1**). *Stat5a* expression was also downregulated in the radial glial cells of the fetal brain in the absence of *Cav1* further supporting a role of *Cav1* in the regulation of MPB axis.

However, we want to point out that our study doesn’t provide evidence for a causal role of mammary gland in gene regulation of placenta and fetal brain. It should be emphasized that the mouse used in this investigation was a *Cav1* global knockout model, not a conditional knockout model specific to the mammary gland. It is our aim in the future direction of this work to generate a conditional knockout mouse model to specifically knockout the *Cav1* gene from the mammary gland and then investigate its effects on placenta and fetal brain development.

Nevertheless, the findings of the present study provide evidence for a role of *Cav1* in the molecular crosstalk between of the mammary gland, placenta and fetal brain. As *Cav1*-null mice exhibit symptoms of neurodegeneration early in adult life (Head et al. 2010), our findings suggest that mammary gland metabolism during pregnancy may be linked to placental function and fetal brain programming. In conclusion, the findings highlight the importance of mammary gland metabolism to pregnancy and fetal development (Gage et al. 2016; Goldstein et al. 2020), and underscore the need for further research to investigate role of mammary gland metabolism in the regulation of fetoplacental communication during pregnancy.

## Supporting information

Metabolites and their level in the mammary gland of control and Cav1-null mice (n=3).

The mean level and rank order of metabolites in the mammary gland of control and Cav1-null mice.

List of metabolic pathway genes expressed in the mammary gland (MG) of control (C) and Cav1 knockout (KO) mice (n=3).

List of metabolic genes showing increased expression in the mammary gland (MG) but decreased expression in the fetal brain and placenta of Cav1-null

List of genes associated with different clusters identified by hierarchical clustering.

List of pathways significantly enriched with the metabolic and signaling genes. The fold and significance of enrichment are shown for each pathway.

The aggregated expression of the metabolic (shown as mDEG), cytokine and cytoskeleton genes in the fetal brain and placenta cells of the control and C

Expression of Stat5a gene in different cell types of the placenta and fetal brain. Dot plots show the expression level and percentage (of cells) that

## Acknowledgements

The authors are thankful to the University of Missouri Metabolomics Center, and also the Genomics Technology core for providing the metabolomics and next-generation sequencing services during this study.

## Conflicts of Interest Statement

The authors declare no conflicts of interest.

## Funding

This study was supported in parts from the AG2PI seed grant, and a start-up fund from the University of Missouri.

## Author Contributions

SKB, TBM conceived and designed research; SPP, IM,TBM performed experiments; SKB, SPP analyzed data; SKB, SPP, TBM wrote the paper. All authors read and approved the manuscript.

## Ethical Statements

All applicable international, national, and/or institutional guidelines for the care and use of animals were followed.

## Data Availability

The raw and processed data of RNA-seq assays have been submitted to the Gene Expression Omnibus (GEO) database under the accession number GSE268570. The raw and processed data of the metabolomics assay are available at Metabolomics Workbench under study ID ST003855 and accession ID SbsM9645.

## Abbreviations

*Cav1*: Caveolin-1;
FDR: false discovery rate;
GO: gene ontology;
GC: Gas chromatography;
MS: Mass spectrometry;
TMCS: N-Trimethylsilyl-N-methyl trifluoroacetamide; trimethylchlorosilane,
AMDIS: Automated Mass-spectral Deconvolution and Identification System;
MET-IDEA: Metabolomics Ion-Based Data Extraction Algorithm;
DE: Differential expression;
MRMR: Maximum relevance minimum redundancy;
MI: Mutual information;
CTB: Cytotrophoblast,
EVT: Extravillious trophoblast,
STB: Syncytiotrophoblast

## Supplementary Data

**Supplementary Figure 1.** Expression of *Stat5a* gene in different cell types of the placenta and fetal brain. Dot plots show the expression level and percentage (of cells) that express *Stat5a* in different cell types (shown in y-axis) of placenta of control mice (**A**), placenta of *Cav1*-null mice (**B**), fetal brain of control mice (**C**) and fetal brain of *Cav1*-null mice (**D**).

**Supplementary Table 1.** Metabolites and their level in the mammary gland of control and *Cav1*-null mice (n=3).

**Supplementary Table 2.** The mean level and rank order of metabolites in the mammary gland of control and *Cav1*-null mice.

**Supplementary Table 3.** List of metabolic pathway genes expressed in the mammary gland (MG) of control (C) and *Cav1* knockout (KO) mice (n=3).

**Supplementary Table 4.** List of metabolic genes showing increased expression in the mammary gland (MG) but decreased expression in the fetal brain and placenta of *Cav1*-null compared to control mice. Fold changes (FC) of expression and significance p-values are shown.

**Supplementary Table 5.** List of genes associated with different clusters identified by hierarchical clustering.

**Supplementary Table 6.** List of pathways significantly enriched with the metabolic and signaling genes. The fold and significance of enrichment are shown for each pathway.

**Supplementary Table 7.** The aggregated expression of the metabolic (shown as mDEG), cytokine and cytoskeleton genes in the fetal brain and placenta cells of the control and *Cav1* knockout mice. CTB: cytotrophoblasts, EVT: extravillious trophoblasts, and STB: syncytiotrophoblasts.

## Notes

https://www.ncbi.nlm.nih.gov/geo/

https://www.metabolomicsworkbench.org/

### Competing Interest Statement

The authors have declared no competing interest.

