## Supplementary figures and images for "Mammary gland metabolism and its relevance to the fetoplacental expression of cytokine signaling in Caveolin-1 null mice"

### Expression of Stat5a gene in different cell types of the placenta and fetal brain. Dot plots show the expression level and percentage (of cells) that

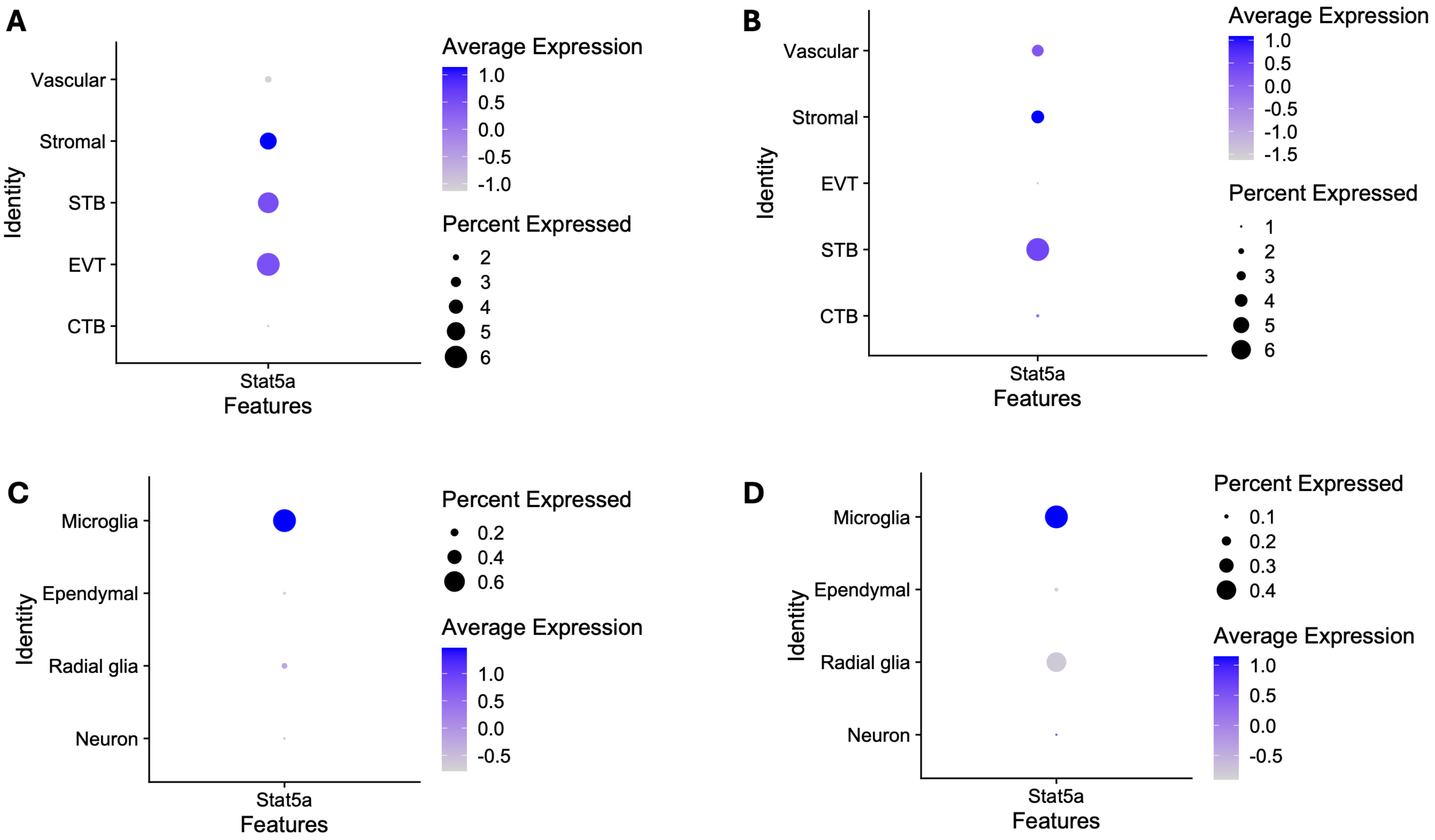
